# Testing the Cross-Paradigm Convergence of Behavioral and Neural Measures of Attention

**DOI:** 10.64898/2026.07.08.737184

**Authors:** Rotem Waxman, Orel Levy, Shun Okada, Elana Zion Golumbic

**Author notes:** joint first authorship (Rotem Waxman, Orel Levy).

## Abstract

Attention abilities, that are critical for most real-life cognitive task, can vary greatly across individuals. A plethora of behavioral and neural metric are commonly utilized to quantify attention; however, these often yield inconsistent and non-replicable results, raising questions as to which measures reliably account for and capture individual differences. To address this tension, here we employed a within-subject cross-paradigm design and quantified both behavioral and EEG-based neural measures associated with attention functioning, to test which measures converged across a gradient of artificial and ecological tasks. We found Inter-Subject Correlation (ISC), which quantifies the similarity in the pattern of neural response across individuals, captured consistent individual differences across both an artificial (Auditory Oddball) and ecological (Attention to Speech) task, with some correlation with performance. Moreover, we found that P300 neural responses to surprising events were qualitatively similar across artificial and ecological tasks, supporting their generalization across contexts. Interestingly, traditional cognitive-behavioral measures of attention – both from speeded response-time tasks and self-report of ADHD symptoms (ASRS) were not correlated with each other nor did they explain variance in neural metrics, nor were they explained by variation in general cognitive abilities (working memory or fluid intelligence). While these results demonstrate the shortcomings of many established measures of attention to generalize across contexts, they also point to the potential of neural measures, and specifically ISC, to serve as reliable indicators of individual differences and to bridge the gap between artificial and ecological studies of attention functioning.

## Introduction

Attention is a critical component of most cognitive processes, including perception, learning, social interaction and decision-making (Chun et al., 2011; Desimone & Duncan, 1995; Petersen & Posner, 2012a; Posner & Petersen, 1990). However, despite its centrality to human behavior, it is not easy to define “what attention is” (Burgoyne et al., 2023; Posner & Boies, 1971). Empirical and clinical studies offer many different operationalization for measuring attentional functioning - running the gamut from self-report questionnaires (Adler et al., 2006; Kessler et al., 2005; Toplak et al., 2013a; Unsworth et al., 2012) through computerized speeded-response tasks (Fan et al., 2002a; Robertson et al., 1997) to more ecological evaluations of attention in real-life settings (Draheim et al., 2022a; Robertson et al., 1996). Complementing behavioral measures, specific neural and physiological measures have been associated with aspects of attentional functioning, often in attempt to dissociate between sub-components of attention such as alerting, sustained attention, orienting and executive control (Fan et al., 2002a; Fortenbaugh et al., 2017; Petersen & Posner, 2012a), or to differentiate between so-called “attentional states” (Dmochowski et al., 2012; Esterman et al., 2013; Kam et al., 2022; Ki et al., 2016a; Klimesch, 2012; Luck et al., 2000; Smallwood & Schooler, 2015).

Although research over the past decades has provided critical mechanistic insight and influential theories regarding the taxonomy, operations and limitations of attention, the plethora of approaches used to quantify “attention” often exhibit mixed results that can be difficult to synthesize into a unified understanding of attention systems (Burgoyne et al., 2023; Friedman & Miyake, 2016; Toplak et al., 2013a). For example, there are increasing reports of inconsistent performance on tasks supposedly designed to target similar underlying constructs and a lack of convergence among neural measures thought to capture “attention abilities” (Duckworth & Kern, 2011; Friedman & Miyake, 2016; Hedge et al., 2017a; Rosenberg et al., 2017; Rouder et al., 2023). In addition, the proposed mechanistic distinction between brain networks supporting different facets of attentional functioning (such as alertness, orienting and executive control) does not necessarily replicate across studies (de Souza Almeida et al., 2021; Fan et al., 2002a, 2009; Ishigami & Klein, 2010a; MacLeod et al., 2010a). Similarly, several neural metrics that have been proposed as ‘biomarkers’ of attention, such as the power of Alpha-band activity or the relative power in specific frequency bands (e.g. theta-beta ratio), do not produce reliable and replicable results (Fortenbaugh et al., 2017; Gloss et al., 2016; Lenartowicz et al., 2018; Loo & Makeig, 2012; Ogrim et al., 2012).

The confusion regarding the optimal means to quantify “attention” is particularly acute in the clinical domain, where “attention deficits” are a clinically-diagnosable condition (American Psychiatric Association, 2022; Musullulu, 2025; Rubia, 2018) and yet the tools used to evaluate attention are highly subjective and vary greatly across clinical practices (Murphy & Adler, 2004; Musullulu, 2025; Kooij et al., 2008). Recent reviews and meta-analyses have pointed out this tension, noting insufficient sensitivity and specificity of many clinical practices for the evaluating attention deficits for clinical purposes (Arrondo et al., 2024a; Barkley, 2019; Chang et al., 2016; Musullulu, 2025) as well as poor reliability of most neural measures for explaining individual-level differences in attentional functioning (Adamou et al., 2020; Clayson et al., 2021; Faraone et al., 2021; Lopez et al., 2023; Noble et al., 2019). One source of these discrepancies is the abundance behavioral tasks and neural metrics that are all supposedly markers of “attention”, but actually vary greatly in their perceptual and cognitive demands, their ecological validity and their reliance of subjective introspection (Draheim et al., 2022a; Friedman & Gustavson, 2022; Smilek et al., 2010; Toplak et al., 2013a; Unsworth et al., 2012). This makes it difficult to separate between variance that is due to task-specific demands vs. true trait-based variability across individuals (Hedge et al., 2017a; Miyake et al., 2000; Rouder et al., 2023).

The basic criterion for a metric – be it behavioral or neural - to serve as a reliable indicator of stable individual-differences in attention, is that it is stable and replicable within-person across different contexts. The current study is an attempt to identify behavioral and/or neural metrics that account for individual variability across different attention-related paradigms, and distinguish them from measures that fail to generalize beyond the specific context. To do so, we employed a within-subject cross-paradigm approach and quantified several different measures associated with attention functioning, that span a gradient of ecological relevance and pose different contextual demands. These included: (1) Behavioral assessments of alertness and executive control using the Attention Network Task (Fan et al., 2002a, 2005), (2) Neural responses associated with selective attention, target detection and attentional capture, using an Auditory Oddball Task (Berti, 2013; Escera & Corral, 2008, 2008; D. Friedman et al., 2001; Polich, 2007, 2007; Schröger & Wolff, 1998), and (3) Behavioral and neural measures on an ecologically-valid attention task, that simulates the attentional demands of paying attention to continuous speech amidst ecological disturbances. Neural measures included stimulus-evoked neural responses, as well as inter-subject correlation (ISC), which quantifies the similarity in neural responses across individuals exposed to the same stimulus and has been linked to engagement with presented stimuli (Dikker et al., 2017; Dmochowski et al., 2012; Hasson et al., 2004; Ki et al., 2016b). We evaluated the correspondence between behavioral and neural measures within and between these tasks, as well as their ability to predict individuals’ introspective self-report of attention functioning using a validated clinical questionnaire (Adult ADHD Self-Report Scale; ASRS; (Adler et al., 2006; Hines et al., 2012; Kessler et al., 2005). Last, we also test whether individual differences in these empirical metrics are linked to personal variations in more general cognitive abilities, such as working memory and generalized intelligence (Diamond, 2012; Engle, 2018; Unsworth et al., 2014). This rich and multimodal data has the potential to broaden our understanding of individual differences in attention beyond the limitations of a single paradigm and to address critical concerns pertaining to generalization and trait-stability.

## Methods

### Participants

Data were collected from 60 adults (33 females, 27 males), aged 18 to 52 years (mean age: 26.35 ± 6.87). All participants self-reported normal hearing and no history of neurological disorders. Five participants had a prior clinical diagnosis of ADHD, and they were included in the sample. Due to technical errors, data from two participants were not available for the computerized ANT task (session 1) and were therefore excluded from analyses involving this task. Participants received either monetary payment or course credit for their participation. The study was approved by the Institutional Review Board at Bar-Ilan University, and all participants provided written informed consent prior to the experiment.

### Experimental Procedure

The experiment consisted of two sessions. Session 1 was conducted online and Session 2 took place in the lab, no more than 10 days after Session 1. Figure 1 illustrates the battery of tasks completed in both sessions.

**Figure 1.**
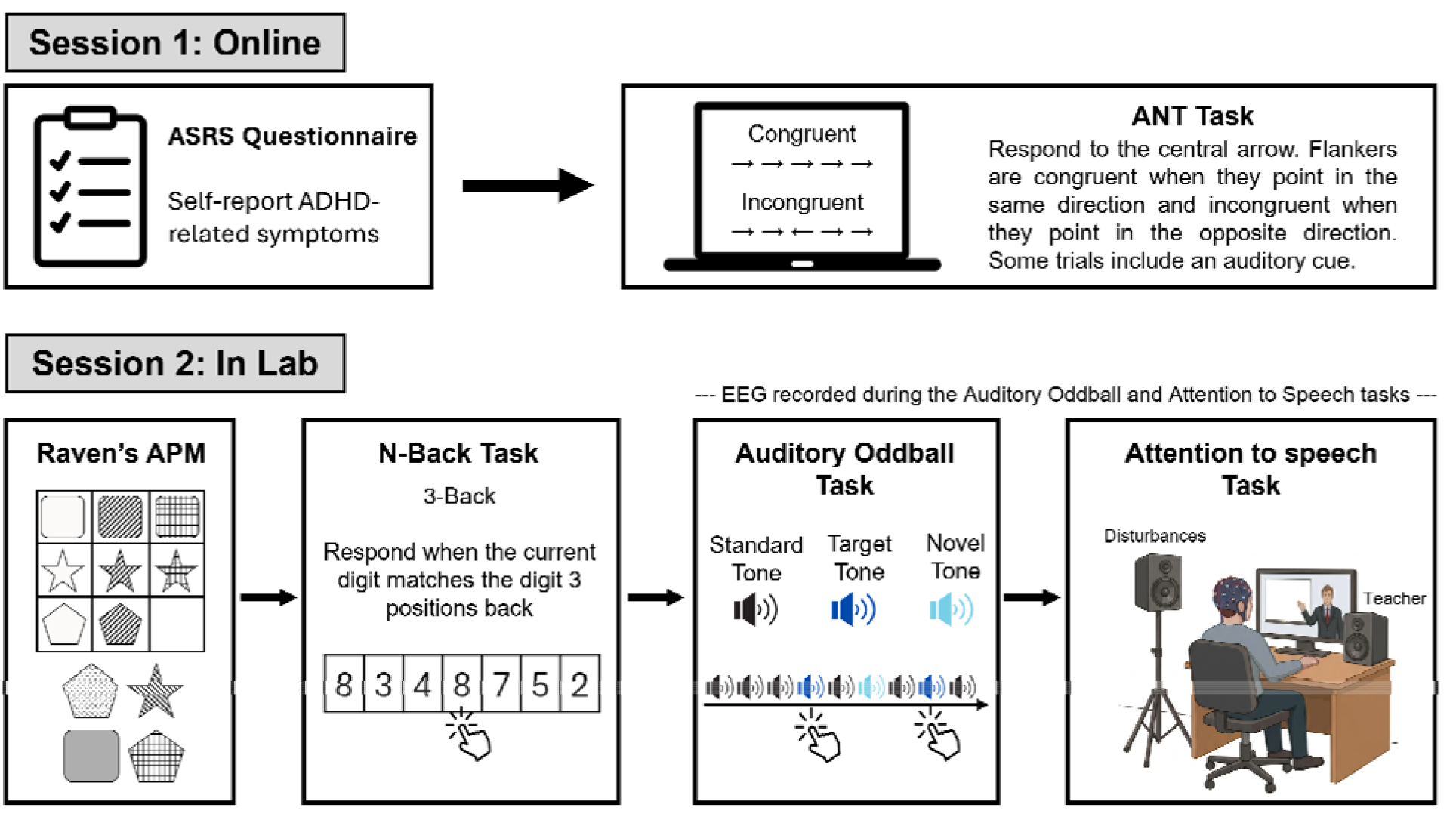
Experimental design. Participants completed two sessions within 10 days. Session 1 was conducted online and included the ASRS questionnaire and the Attention Network Test (ANT). Session 2 was conducted in the laboratory and included Raven’s Advanced Progressive Matrices, an N-back task, an Auditory Oddball task, and an Attention to Speech task. EEG was recorded during the auditory oddball and Attention to Speech tasks.

#### Session 1: Online

The first session was conducted during a live Zoom-call between the experimenter and each participant. Participants were instructed to find a quiet, distraction-free environment with a stable internet connection to complete the session. Participants were asked to keep their webcam and microphone on throughout the session, to allow monitoring and communication with the researcher, as needed. This online session was part of a larger study (N=156), and the full results are reported in Oshri et al. (*in prep)*. Here we report results only from a subset of N=60 participants who also completed the in-lab session 2. During the online session, participants completed an ADHD Self Report Scale (ASRS) questionnaire and performed a computerized Attention Network Task (ANT), as described below:

##### ADHD Self Report Scale (ASRS)

Participants completed an 18-item ASRS questionnaire, a validated and widely used tool for assessing symptoms of Attention Deficits and Hyperactivity Disorder (ADHD), in line with DSM-5 diagnostic criteria (ASRS-v1.1; Adler et al., 2006, Hebrew version: Zohar and Konfortes, 2010), administered using the Qualtrics platform (Provo, UT, https://www.qualtrics.com/) . ASRS scores were computed by summing all item responses, yielding scores ranging from 0 to 72. Based on prior literature, individuals with scores above 51 are considered “candidates for ADHD”. Beside completing this questionnaire, participants were also asked if they had been were previously diagnosed with ADHD.

##### Modified Attention Network Task (ANT)

Participants completed a modified version of the Attention Network Task (ANT) (Reinholdt-Dunne et al., 2009; Roberts et al., 2006; Russman Block et al., 2020; Stone et al., 2021). The original ANT task, as described by Fan et al. (2002), was designed to dissociate between behavioral effects associated with the alertness, orienting and executive-control attentional networks Petersen and Posner (2012). The ANT has been used extensively in attention research (Block et al., 2020; Reinholdt-Dunne et al., 2009; Stone et al., 2021), both online and in lab experiments (yielding comparable results; Luna et al., 2018). Here we used a modified version of ANT, that focuses on two aspects of attention: *Alertness* and *Executive Control* (illustrated in Figure 1).

Each trial started with a 650 millisecond-long fixation point, followed by five horizontal arrows presented for a maximum of 1 second (until button press). Participants were required to report the direction of the **central** arrow using corresponding keyboard keys, irrespective of the direction of surrounding arrows (flankers). To manipulate *Executive Control* demands, in 50% of the trials the flanking arrows faced the same direction as the central arrow (Congruent condition) and in 50% they faced the opposite direction (Incongruent condition). To manipulate *Alertness*, 25% of trials include an auditory cue (440 Hz; 100 ms long) presented 150 ms prior to the appearance of the arrow stimuli (500 ms after fixation onset). Auditory cues occurred with equal frequency in Congruent and Incongruent trials. Trials were separated by a 1-second long blank screen. Participants were instructed to respond as quickly as possible and to also maintain high accuracy. Participants first completed a short training session comprised of 16 trials (from all conditions) and then proceeded to perform the full experiment, which included 256 trials. The task was programmed in PsychoPy version 2020.1 and executed online through the Pavlovia platform (https://pavlovia.org). One limitation of the on-line version is the lack of control over the sound-volume of the tones. To address this, prior to the start of the experiment participants were presented with the auditory stimuli used in the experiment and were asked to adjust the volume of their headphones so that they could *“hear the sound clearly”*. Admittedly, this is not akin to a careful calibration process, and therefore it is possible that some variance between participants is due to different perceived levels of loudness of the auditory stimuli. Analysis of behavioral responses focused on accuracy and reaction-times (RTs). Following the approach of Fan et al. (2002), Alertness effects were assessed by calculating the difference in mean RT between trials with and without an auditory cue (Cue − No cue), and Executive Control effects were assessed by calculating the difference in mean RT between Congruent and Incongruent trials (Congruent − Incongruent).

#### Session 2: In-Lab

The second session was conducted in the lab at Bar Ilan University. It consisted of four tasks: Two Behavioral-Cognitive tasks (Raven’s Advanced Progressive Matrices and N-Back working-memory task) and two EEG tasks (an Auditory Oddball task and an Attention to Speech task).

##### Raven’s Advanced Progressive Matrices (Behavioral Task)

This is a non-verbal test that is widely used to assess general cognitive ability (“fluid intelligence”) (Carpenter et al., 1990). It involves identifying perceptual relations and reasoning by analogy to solve novel visuo-spatial problems (Raven et al., 1998; Cattell, 1963). Here we used a shortened 12-item version, validated by (Bors & Stokes, 1998) The task was programmed in PsychoPy version 2020.1 and executed online through the Pavlovia platform (https://pavlovia.org).

##### N-Back (Behavioral Task)

This task is widely used to assess Working-Memory (WM) and continuously internal-model updating (Ackerman et al., 2005; Kirchner, 1958; Owen et al., 2005). Here we used a 3-back version of the task, where participants viewed a sequence of 70 digits and were instructed to respond via mouse click whenever the current digit matched the one presented *three trials earlier*. Each stimulus was preceded by a fixation cross presented for a randomly jittered interval ranging from 0.5 to 1.5 seconds, in order to reduce temporal predictability and minimize anticipatory responses. The task was programmed in PsychoPy version 2020.1 and executed online through the Pavlovia platform (https://pavlovia.org).

##### Auditory Oddball (EEG Task)

This task is designed to measure neural event-related potentials (ERPs) associated with sensory and attention-related responses to different sounds (Squires et al., 1975). Specifically, we used a 3-stimulus version that contrasts responses to Standards (1000 Hz pure tones), Targets (1500 Hz pure tones; requiring a button-press response), and ecological Novel sounds (e.g., phone ringing, glass breaking) (Masson & Bidet-Caulet, 2019; O’Connor et al., 1994). Standard and Target tones were 50ms in duration with a 10ms ramp-up and ramp-down, created using Audacity software. Novel sounds were 300ms long and taken from (Masson & Bidet-Caulet, 2019). Each experimental block consisted of 80 sounds with a distribution of 70% Standards, 15% Targets, and 15% Novels. Stimulus order was pseudo-randomized under two constraints: (1) a minimum of five Standard tones at the beginning and end of each block, and (2) at least one Standard tone separating any Target and Novel. The inter-stimulus interval (ISI) was fixed at 1000 ms, making each block approximately 87 seconds long. The task was programmed and presented using OpenSesame software (https://osdoc.cogsci.nl). Auditory stimuli were presented free-field at 70dB through a centrally positioned loudspeaker placed behind the computer screen in front of the participants.

##### Attention-to-Speech (EEG task)

This task was designed to simulate the type of attention demands posed under realistic condition, and simulates the experience of video-based learning (Cohen et al., 2018a; Davidesco et al., 2023; Levy, Shadi, et al., 2025). Participants watched a ∼20-minute video lecture presented on a computer monitor in front of them. The topic of the lecture was Savant syndrome, and was delivered by a professional lecturer and optimized for on-line learning. The lecture was divided into 12 consecutive segments (98.7 ± 16.8 seconds each) each followed by three multiple-choice questions used to assess comprehension of the preceding content. In addition, half of the segments included background speech disruptions that mimic the experience of a nearby student making casual remarks. These included short utterances such as (e.g., “*this is so interesting,” “it might rain,” “I’m so tired”*; 2.22 ± 0.43 seconds each). Notably, each disturbance was preceded by the Hebrew interjection “Wai” (analogous to “Aw” or “Oh” in English) to provide a naturalistic and acoustically consistent onset for all stimuli. Six disruption events occurred per segment (ISI: 11.55 ± 0.49 s, range: 10.75–12.33 s), yielding 36 events in total. The assignment of specific lecture-segment to condition (Quiet vs. with Disruptions) was counterbalanced across participants. Lecture audio was delivered through a loudspeaker positioned centrally behind the monitor and disruptions were presented through a second loudspeaker positioned to the rear-left of the participant (both placed 95cm from the participant), mimicking the spatial separation experienced in real-world environments.

### EEG acquisition and analysis

Electroencephalography (EEG) was recorded during the Oddball and Attention to Speech tasks using a 64 Active-Two system (BioSemi; sampling rate: 2048 Hz) with Ag-AgCl electrodes. Two external electrodes were placed on the mastoids and served as reference channels. Electrooculographic (EOG) signals were simultaneously measured by 4 additional electrodes, located above and beneath the right eye and on the external side of both eyes. EEG data was synchronized to the auditoy stimuli and digital triggers using Lab Streaming Layer (LSL;Kothe et al., 2025, https://github.com/sccn/labstreaminglayer). EEG data preprocessed using the MATLAB-based FieldTrip toolbox (Oostenveld et al., 2011; version: 20220729; https://www.fieldtriptoolbox.org) and custom-written scripts. Data preprocessing included re-referencing to the linked left and right mastoids, detrending, demeaning, and band-pass filtering the data between 0.5 and 40 Hz using a zero-phase, two-pass Butterworth IIR filter. Artifacts were removed through visual inspection and independent component analysis (ICA), with EOG channels included in the decomposition to facilitate the identification of ocular artifacts. Excessively noisy electrodes were temporarily excluded from ICA to improve decomposition stability and were subsequently reinstated and interpolated using spatially neighboring electrodes. Finally, the data were visually inspected, and remaining non-ocular artifacts were flagged and removed from subsequent analyses.

#### EEG Analysis: Auditory Oddball Task

Continuous EEG data were segmented into epochs time-locked to stimulus onset (-100ms to +900ms) separately for each stimulus type (Standard, Target and Novel). Epochs were visually inspected for residual artifacts and rejected if they exhibited extremely large amplitudes (>5 µV, threshold determined based on the distribution of RMS across all channels and time). Clean single trials were then bandpass filtered (1–20 Hz) and baseline corrected relative to the pre-stimulus period (-100ms to 0).

For each participant and stimulus type we computed two metrics: First, we calculated the **Event-Related Potentials (ERPs)** that capture the average response to each stimulus, across all trials. Event-related potentials (ERPs) were computed by averaging trials within each condition (Standard, Target, and Novel), and statistical analysis focuses on electrodes Fz and Cz where these responses were maximal. ERPs exhibited three primary and canonical components – The early N1, and later P3 and N4 responses. Fixed time windows were defined for each component based on the grand-average waveforms [N1: 90–180 ms; P3: 190–350 ms; N4: 380–490 ms] and were applied uniformly across all participants and conditions. Component magnitude was quantified for each participant and condition as the mean amplitude within the corresponding time window. We tested for statistical differences in ERP amplitudes between conditions using a one-way repeated-measures analysis of variance (ANOVA) with Condition (Standard, Target, Novel) as a within-subject factor. When a significant main effect was observed, pairwise comparisons between conditions were conducted using Bonferroni-corrected paired-samples t-tests.

The second metric evaluated was the **ERP inter-subject correlation (ERP-ISC;** also referred to as “typicality”). This measure quantifies the similarity between each participant’s ERP response and the group response. Specifically, for each participant and condition, ERP waveforms were first averaged across electrodes Fz and Cz. Then pairwise Pearson correlations were calculated between that participant’s ERP waveform and the ERPs of all other participants in the same condition. These pairwise correlations were then averaged to yield a single ERP-ISC value per participant in each condition (see Korisky et al., 2026). To test whether ERP-ISC differed across stimulus types, values were compared using a one-way repeated-measures ANOVA with Condition as a within-subject factor. Follow-up pairwise comparisons were conducted using Bonferroni-corrected paired-samples t-tests.

To estimate the proportion of ERP-ISC variability attributable to participants, task conditions, and residual error, we performed a complementary mixed-effects intraclass correlation analysis (ICC). ERP-ISC values were modeled using a linear mixed-effects model with random intercepts for participant and condition: ERP-ISC ∼ (1 | Participant) + (1 | Condition). Variance components were extracted from the fitted model and divided by the total variance, yielding the proportion of variance attributable to between-participant differences, between-condition differences, and residual variability. The significance of the participant and condition random effects was assessed using likelihood-ratio tests, implemented in the ranova function from the lmerTest package in R.

#### EEG analysis: Attention to Speech Task

For the Attention to Speech task we report two types of analysis – Event Related and Continuous: For the event-related analysis of responses to disruption onsets (the “Wai” stimulus) we calculated both ERPs and ERP-ISC, using the same procedures described for the Auditory Oddball task (above). Throughout the paper, we will refer to the latter as **Disruption-ISC**. In addition, we conducted **Continuous-ISC** analysis of the ongoing neural activity while watching the lecture. This measure is particularly suitable for analyzing neural response to continuous, dynamic stimuli and quantifies the degree of similarity in participant’s neural-dynamics over time which has been proposed as a useful metric of attention and/or engagement with continuous stimuli (Cohen & Parra, 2016; Ki et al., 2016a; Poulsen et al., 2017). To calculate the continuous-ISC, EEG data from the full-length lecture segments (98.7 ± 16.8 s each) were analyzed using the open-source Parra lab toolbox (https://www.parralab.org/isc/) to perform correlated component analysis (CorrCA), which estimates spatial filters that maximize between-subject covariance relative to within-subject covariance (Parra et al., 2019). Each participant’s multichannel EEG was projected onto these filters, yielding a time series for each subject per component. Subject-level ISC was then calculated separately for each component by quantifying the covariance between each participant’s projected response and the projected responses of the other participants, normalized by the corresponding within-subject covariance, following the procedure described by Cohen & Parra (2016). Based on previous studies the continuous ISC values were defined as the sum of all pairwise correlations for the first three correlated components (Dmochowski et al., 2012; Ki et al., 2016a; Petroni et al., 2018; Rosenkranz et al., 2021). The scalp projections of the first three components used to compute the ISC summary measure are shown in Figure 5.

#### Statistical analyses

Statistical analyses were conducted in Python using pandas for data handling and statistical packages, such as SciPy, statsmodels, and Pingouin, as we as the lmerTest package in R.

## Results

### Cognitive-Behavioral Tasks

Three cognitive-behavioral tasks were included in this study: ANT, Raven APM tasks and N-back. We first provide descriptive statistics for each task separately, and then report the correlations between performance metrics across tasks (Figure 2):

**Figure 2.**
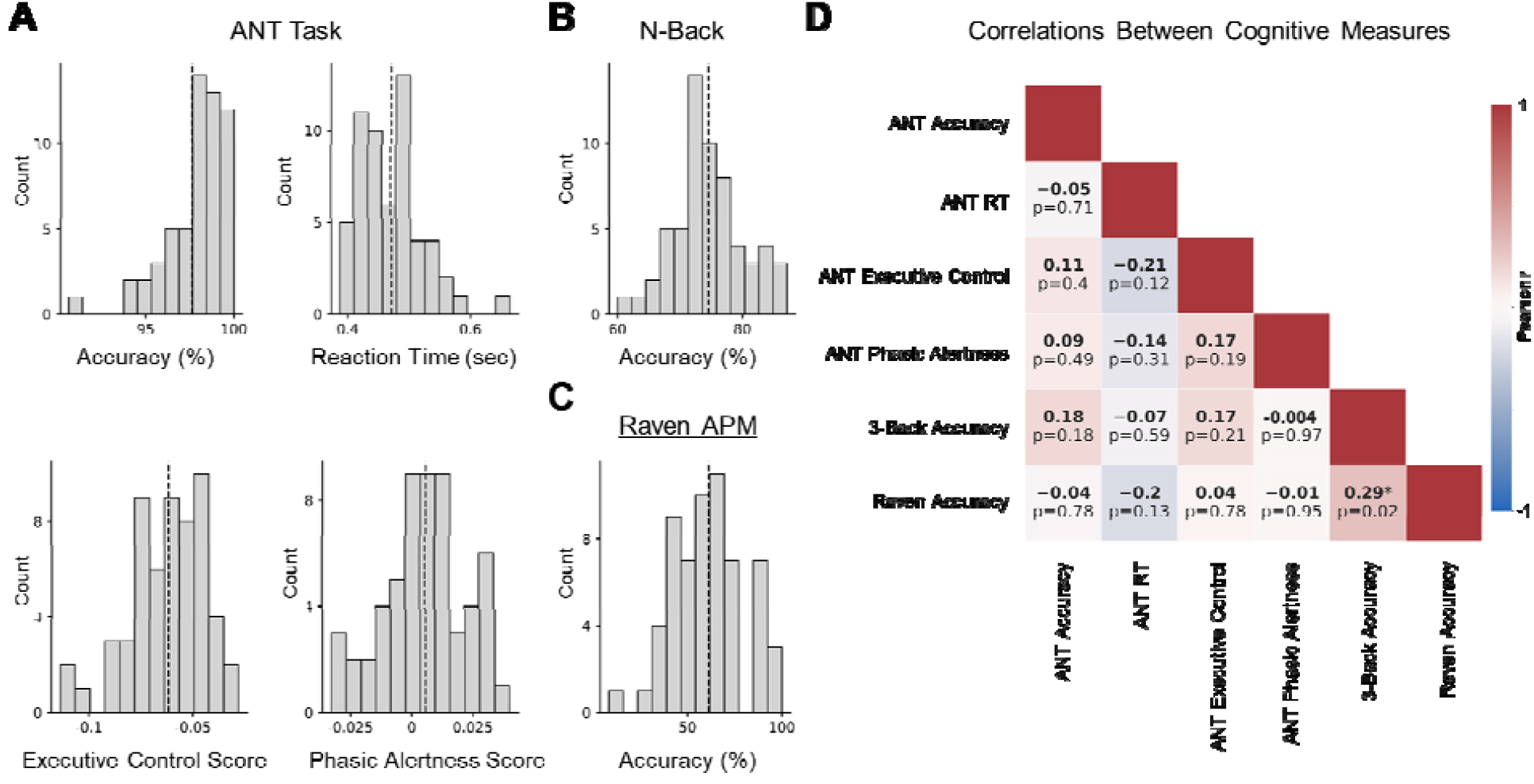
Cognitive-Behavioral tasks results. (A) Distribution of the four performance measures derived from the ANT task. Participants showed high accuracy (M = 97.64%, SD = 1.79) and significant effects of executive control (RT difference between Congruent vs. Incongruent trials) and phasic alertness (RT difference between trials with vs. without an auditory cue), although effect sizes were small. (B) Distribution of accuracy on the N-back task (M = 74.57%, SD = 5.83). (C) Distribution of accuracy on the Raven APM task (M = 61.25%, SD = 20.29). (D) Pairwise Pearson correlations between behavioral measures with and between tasks. The only nominally significant association was between Raven APM accuracy and N-back accuracy but this correlation did not survive correction for multiple comparisons (r = 0.29, p = 0.025, FDR corrected p = 0.37).

#### ANT Task

Participants performed this task with high accuracy overall (M = 97.635%, SD = 1.793%) and a mean reaction time of 471.298 ms (SD = 50.541 ms). Reaction times showed the expected Congruency effect, with slower responses on Incongruent relative to Congruent trials [t(56) = -26.40, p < .001], with substantial variability in the size of the Congruency effect across individuals (range = 28–114 ms, SD = 17.50 ms). Participants also showed a phasic alertness effect, with slightly faster responses following alerting cues [t(56) = 2.50, p = .016]. However, here too the magnitude and direction of this effect varied across participants, and in some cases was observed in the opposite direction (range = -32.85 to 39.90 ms, SD = 16.84 ms; Figure 2A).

#### N-Back WM task

Performance on the N-back WM task was moderately good, with a mean accuracy rate of 74.57% (SD = 5.83), indicating that the task was indeed challenging. Scores were roughly normally distributed with no extreme outliers (Figure 2B).

#### Raven APM task

Performance on the Raven task also exhibited substantial variability across participants, with an average accuracy of 61.4%, and large variability across participants (SD = 20.46; range: 8.33-100%; Figure 2C).

#### Correlation between Cognitive-Behavioral Tasks

Figure 2D shows the pairwise correlations between the performance measures on all three tasks. The ANT task yielded four different behavioral measures – Accuracy, RT, Executive Control effect, and Phasic alertness effect – none of which were correlated with each other (all ps > .05, uncorrected). In addition, none of the ANT-based metrics were significantly correlated with performance on the Raven or N-back tasks (all ps > .05). Accuracy on the Raven and N-back tasks did show a moderate positive correlation however, this effect did not survive correction for multiple comparisons (r = 0.290, uncorrected p = .025, FDR corrected p= .370). Taken together, these results do not indicate a convergence among the different behavioral measures of cognitive/attention-related abilities tested here.

### EEG Tasks

#### Auditory Oddball task

Behavioral performance on the Oddball task was near ceiling, with a mean hit rate of 97.87% (SD = 3.68; range = 80.21–100), indicating good task understanding and compliance. Neural ERPs showed the canonical response patterns typically associated with the three stimulus types (Standard, Target, and Novel; Figure 3A&B). All three stimulus types elicited an early N1 response (∼72 ms), with the largest amplitudes observed for Novel sounds and the smallest for Standard sounds. A one-way repeated-measures ANOVA confirmed a significant main effect of Condition on N1 amplitude [F(2, 118) = 66.51, p < .001, η^2^ = .530]. Follow-up Bonferroni-corrected paired-samples t-tests indicated that N1 amplitudes were significantly larger for Target than Standard stimuli [t(59) = 5.51, p < .001], and significantly larger for Novel than both Standard stimuli [t(59) = 10.10, p < .001], and Target stimuli [t(59) = 6.95, p < .001]. Standard sounds also elicited a P2 response, peaking at ∼190ms, whereas Novel sounds elicited a large P300 response around 290ms, followed by a negative peak around 400ms, consistent with a N400 (at ∼401ms).

**Figure 3.**
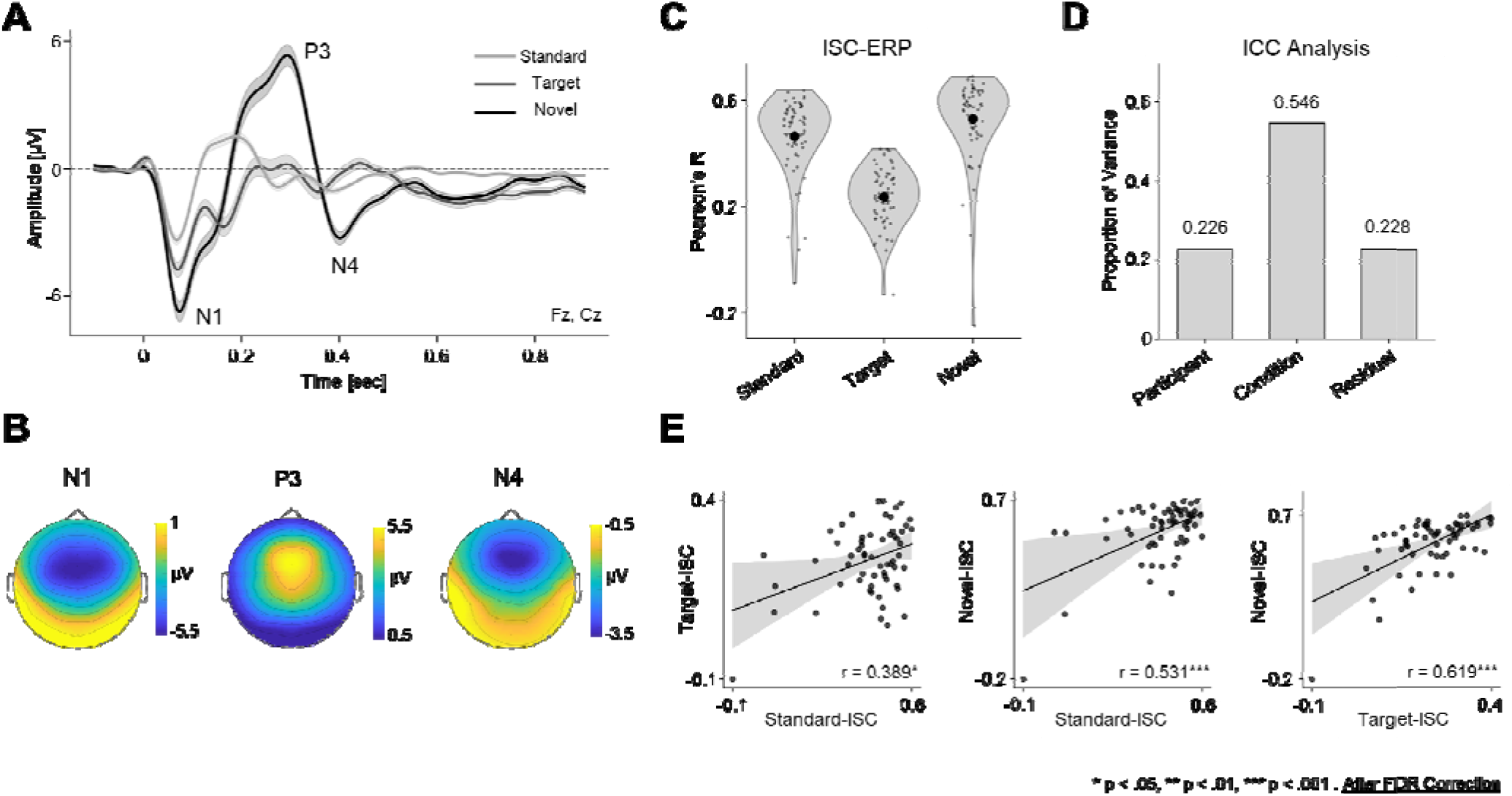
Auditory Oddball Results. (A) ERP waveforms for Standard, Target, and Novel sounds averaged across electrodes Fz and Cz. Novel sounds elicited the strongest response, including a pronounced N1, P3 and N4. (B) Scalp topographies of the three main components in the ERP to Novel sounds. (C ERP-ISC values for Standard, Target, and Novel sounds, showing lower consistency between participants for Target stimuli relative to the other two. (D) Variance decomposition from the intraclass correlation analysis (ICC), showing that stimulus condition accounted for the largest proportion of variance, with additional variance attributable to between-participant differences. (E) Pairwise correlations between ERP-ISC values across conditions, showing strong positive associations among Standard, Target, and Novel ERP-ISC.

ERP-ISC also differed significantly across stimulus conditions, as confirmed by a one-way repeated-measures ANOVA [F(2, 118) = 144.67, p < .001, η^2^⍰ = .710; Figure 3C]. Bonferroni-corrected paired-samples t-tests revealed higher ERP-ISC for Novel sounds than for both Standard sounds [t(59) = 3.35, p = .004], and Target sounds [t(59) = 18.19, p < .001], as well as higher ERP-ISC for Standard than Target sounds [t(59) = 12.15, p < .001]. Nevertheless, ERP-ISC values were significantly correlated among all stimulus types, indicating partial stability of individual differences across conditions (Standard–Target: r = .40, p < .01; Standard–Novel: r = .54, p < .001; Target–Novel: r = .62, p < .001; Figure 3E). This was confirmed in a complementary mixed-effects inter-class correlation (ICC) analysis, showing that 54.6% of the variance in Oddball-ISC was attributable to condition-level differences, 22.6% to between-participant differences, and 22.8% to residual variance (Figure 3D). Both random effects were significant (both p’s < .001).

#### Attention to Speech Task

Accuracy during the lecture task was generally high (M = 83.91%, SD = 8.67), yet varied substantially across individuals, with performance ranging from just above 58% to over 94%. Analysis of neural responses in the semi-naturalistic paradigm focused separately on event-related response to disruption-onset (“Wai” stimulus) as well as analysis of continuous-ISC while listening to the lecture. As shown in Figure 4A&B, the waveform and scalp-distribution of the grand-average ERP to the “Wai” disruption shares many features with the ERP observed for Novel sounds in the oddball paradigm. Specifically, there is a clear negative N1 component peaking around 1210ms (110–133ms), followed by a pronounced positive complex spanning from ∼190–332 ms, which is reminiscent of a P300 response (note: the ‘double-hump’ morphology of this component suggests the possibility that it is comprised of two subcomponents, akin to the P3a and P3b, however the topographical distribution of both peaks was identical, so this remains unresolved). In addition, a negative N400-like response is observed between 423–464ms, peaking at 446ms. We also computed the ERP-ISC in response to the “Wai” disruption, as done for the ERPs in the Oddball paradigm, as well as the continuous ISC. Interestingly, there was a significant correlation between the ERP-ISC to disturbances and the continuous ISC (r = 0.380, uncorrected p = .003, FDR corrected p = .009; Figure 4D&E). Continuous ISC was also significantly correlated with accuracy on answering comprehension questions (r = 0.275, uncorrected p = .035, FDR corrected p = .053), supporting its association with joint attention. In contrast, no significant correlation was found between accuracy and the ERP-ISC to disturbances (r = 0.154, n.s.).

**Figure 4.**
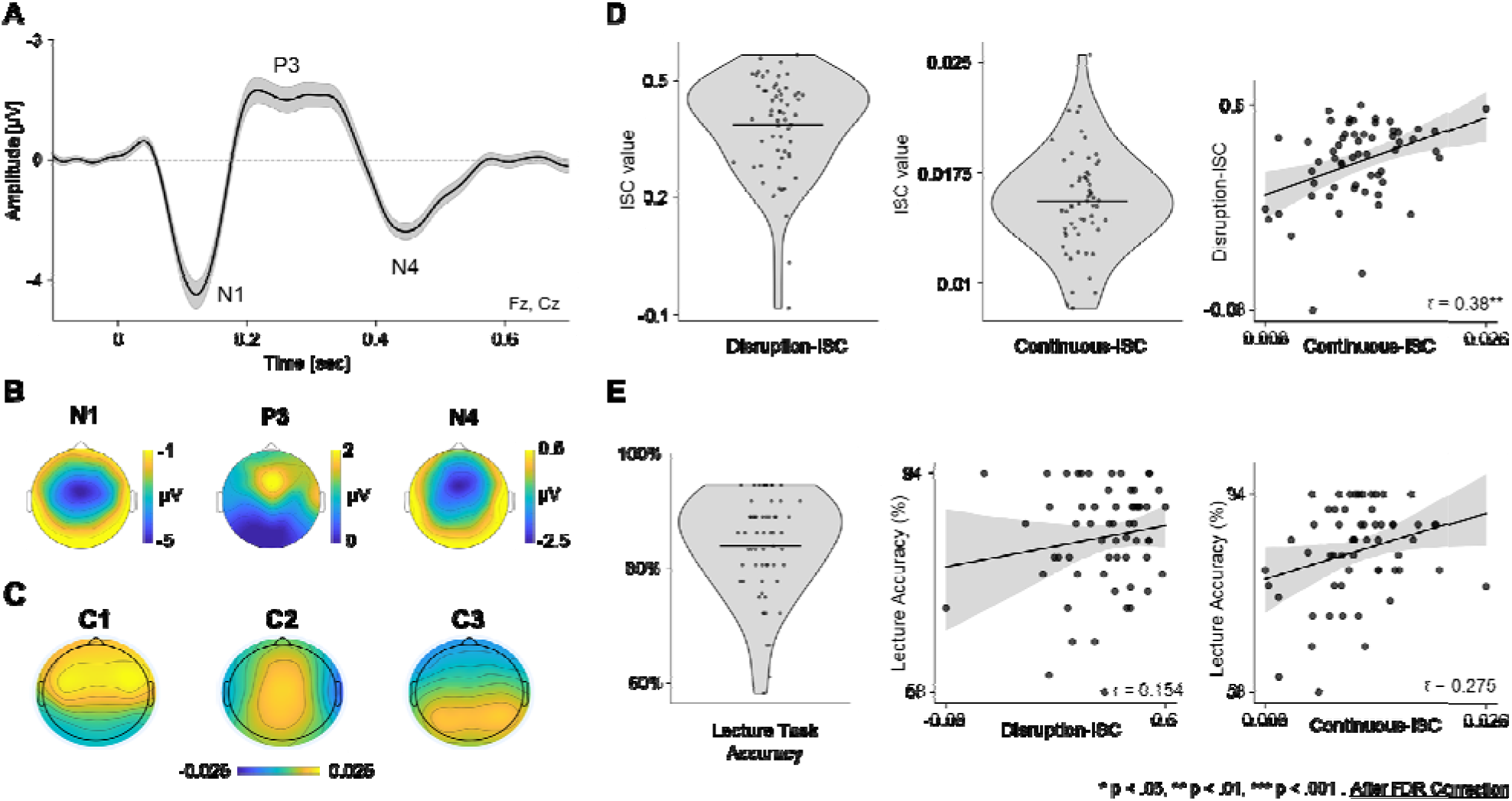
Attention to Speech Results. (A) ERP waveform time-locked to auditory disturbance onset, averaged across electodes Fz and Cz. Disturbance sounds elicited a pronounced N1, followed by a P3 and a later N4 component. (B) Scalp topographies of the three main components in the disturbance-elicited ERP (C) Scalp projections of the first three CorrCA components used to compute Lecture-ISC. (D) Distributions of the ERP-ISC values to the Disruptions (Disruption-ISC; left) and the continuous ISC throughout the entire lecture (middle), as well as the correlation between them (right), which shows a strong positive association. (E) Distribution of accuracy values on the tasks and its correlation with Disruption-ISC and Continuous-ISC values (middle and right). Accuracy was positively association with Continuous-ISC. * p < .05, ** p < .01, *** p < .001, FDR Corrected.

### Comparison of Neural measures in the Oddball vs. Attention to Speech Tasks

As reported above, ISC measures derived separately *within* each paradigm were correlated with each other (ERP-ISC to the three stimuli in the Oddball paradigm were correlated amongst themselves and Disturbance-ISC in the Attention to Speech task were correlated with Continuous-ISC). Here we tested whether these values were correlated *across tasks* (Figure 5). We found that Disruption-ISC was e strongly positively correlated with the ERP-ISC to all three stimuli from the Oddball paradigm, the strongest correlation being with Novel and Target sounds (Novel: r = .346, uncorrected p = .007, FDR corrected p = .019; Target: r = .330, uncorrected p = .01, FDR corrected p = .022), and a weaker correlation with Standard sounds (that did not survive multiple comparisons: r = .26, uncorrected p = .044, FDR corrected p = .067). In addition, Continuous-ISC in the Attention to Speech task was positively correlated with the ERP-ISC to Novel sounds in the Oddball paradigm (r = .322, p = .013, FDR corrected p = .024), but not with Target or Standard sounds (r = .2, p = .128, n.s.; r = .176, p = .183, n.s., respectively).

**Figure 5.**
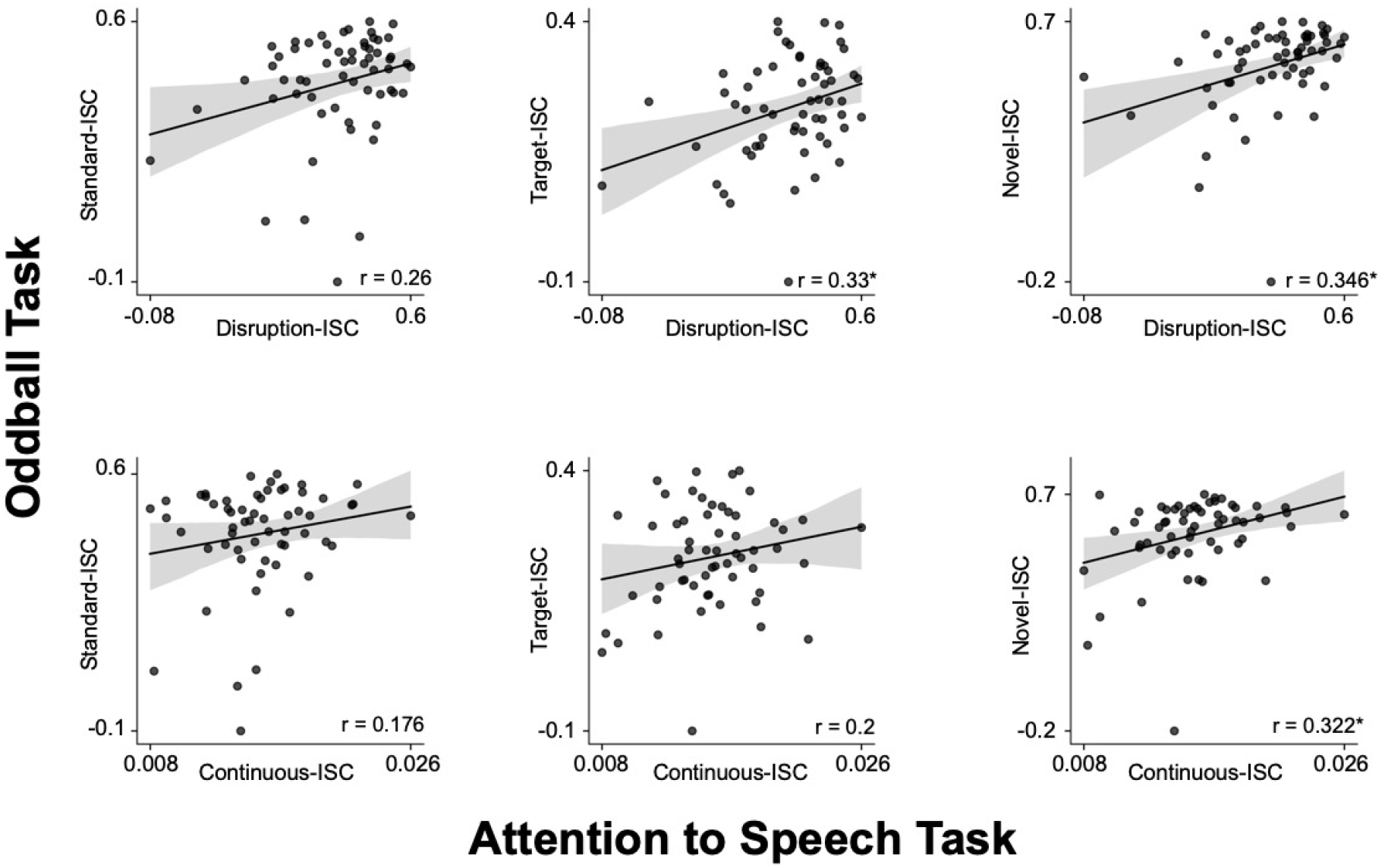
Correlations between ISC measure across tasks. Scatter plots show correlations between ISC to the Standard, Target & Novel stimuli in the Oddball Task vs. the Disruption-ISC, and Continuous-ISC measures from the Attention to Speech task. * p < .05, ** p < .01, *** p < .001, FDR

### Correlation between neural ISC and Performance on Cognitive-Behavioral Tasks

We also tested whether ISC measures from the Auditory Oddball and Attention to Speech paradigms were associated with individual differences in performance on the various cognitive-behavioral tasks. As shown in Table 1, most correlations were non-significant. However, we did find that Novel ERP-ISC from the Oddball task was significantly correlated with comprehension accuracy on the Attention to Speech tasks [r = .341, p = .008, FDR corrected p = .019], an effect that aligns with the correlation between neural metrics reported above.

**Table 1.** Correlations between performance on the cognitive-behavioral tasks and neural ISC measures. Pearson’s correlation coefficient (r), the corresponding uncorrected p-value, and the FDR-corrected p-value. Bold values denote correlations with p < .05 before correction, and correlations that remained significant after FDR correction are further denoted with an asterisk *.

|  |  | ANT<br>Accuracy | ANT<br>Reaction<br>time | ANT<br>Phasic<br>Alertness | ANT<br>Executive<br>Control | Raven<br>Accuracy | N-Back<br>Accuracy | Lecture<br>Accuracy |
| --- | --- | --- | --- | --- | --- | --- | --- | --- |
| Disruption-ISC | $r$ | 0.056 | 0.09 | 0.235 | -0.039 | 0.059 | 0.011 | 0.154 |
| | $p$ | 0.678 | 0.505 | 0.079 | 0.771 | 0.657 | 0.936 | 0.241 |
| | $p$-FDR | 0.912 | 0.891 | 0.393 | 0.963 | 0.912 | 0.969 | 0.648 |
| Lecture-ISC | $r$ | -0.122 | 0.032 | 0.143 | -0.003 | 0.109 | -0.074 | 0.275 |
| | $p$ | 0.369 | 0.814 | 0.292 | 0.98 | 0.413 | 0.577 | **0.035** |
| | $p$-FDR | 0.851 | 0.969 | 0.796 | 0.98 | 0.872 | 0.899 | 0.27 |
| Oddball-ISC<br>(Standard) | $r$ | -0.302 | 0.052 | 0.257 | -0.017 | -0.124 | 0.194 | -0.026 |
| | $p$ | **0.022** | 0.699 | 0.054 | 0.868 | 0.347 | 0.137 | 0.844 |
| | $p$-FDR | 0.324 | 0.912 | 0.324 | 0.969 | 0.851 | 0.586 | 0.956 |
| Oddball-ISC<br>(Target) | $r$ | -0.011 | 0.026 | 0.277 | 0.071 | 0.141 | 0.251 | 0.196 |
| | $p$ | 0.932 | 0.847 | **0.037** | 0.599 | 0.284 | 0.053 | 0.133 |
| | $p$-FDR | 0.969 | 0.969 | 0.324 | 0.899 | 0.796 | 0.324 | 0.479 |
| Oddball-ISC (Novel) | $r$ | -0.19 | 0.074 | 0.158 | 0.096 | 0.102 | 0.256 | **0.341** |
| | $p$ | 0.156 | 0.585 | 0.24 | 0.477 | 0.436 | **0.048** | **0.008** |
| | $p$-FDR | 0.586 | 0.899 | 0.796 | 0.891 | 0.872 | 0.324 | **0.019\*** |

### Correlation with self-report of attention deficit symptoms (ASRS scores)

Last, we looked at the ASRS scores of participants, that are considered indicator for the severity of symptoms associated with ADHD, and are routinely used for clinical evaluation. ASRS scores were widely distributed across participants (Figure 6), with a mean of 50 (SD = 13.21). Notably, a score above 51 is the accepted threshold for being considered “at risk for ADHD”, which here corresponded to the mean and median of the sample. In the current sample, only five participants had a prior clinical diagnosis of ADHD, and although their ASRS scores mostly fell in the upper half of the distribution, they did not stand out by having the most extreme values. Importantly, ASRS scores were not significantly correlated with any of the measures collected here (Table 2, all uncorrected *ps* > 0.05), pointing to a discrepancy between this introspective self-report measure and the more objective behavioral and neural measures measured here.

**Table 2.**
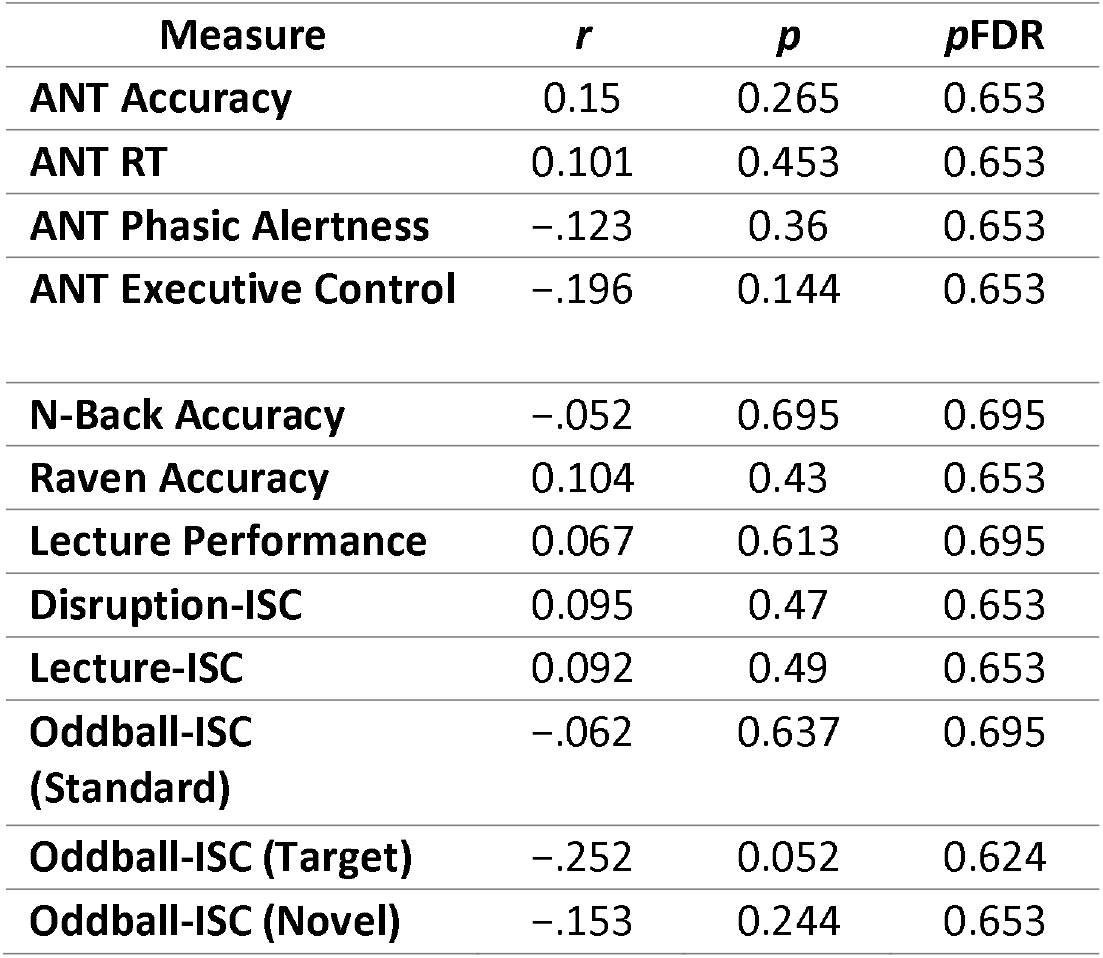
Pearson correlations between ASRS scores and study measures. Pearson correlation coefficients (r), uncorrected p-values, and FDR-corrected p-values are reported for the associations between ASRS scores and the study measures. None of the correlations were significant.

| Measure | $r$ | $p$ | $p$FDR |
| --- | --- | --- | --- |
| ANT Accuracy | 0.15 | 0.265 | 0.653 |
| ANT RT | 0.101 | 0.453 | 0.653 |
| ANT Phasic Alertness | -.123 | 0.36 | 0.653 |
| ANT Executive Control | -.196 | 0.144 | 0.653 |
| N-Back Accuracy | -.052 | 0.695 | 0.695 |
| Raven Accuracy | 0.104 | 0.43 | 0.653 |
| Lecture Performance | 0.067 | 0.613 | 0.695 |
| Disruption-ISC | 0.095 | 0.47 | 0.653 |
| Lecture-ISC | 0.092 | 0.49 | 0.653 |
| Oddball-ISC<br>(Standard) | -.062 | 0.637 | 0.695 |
| Oddball-ISC (Target) | -.252 | 0.052 | 0.624 |
| Oddball-ISC (Novel) | -.153 | 0.244 | 0.653 |

**Figure 6.**
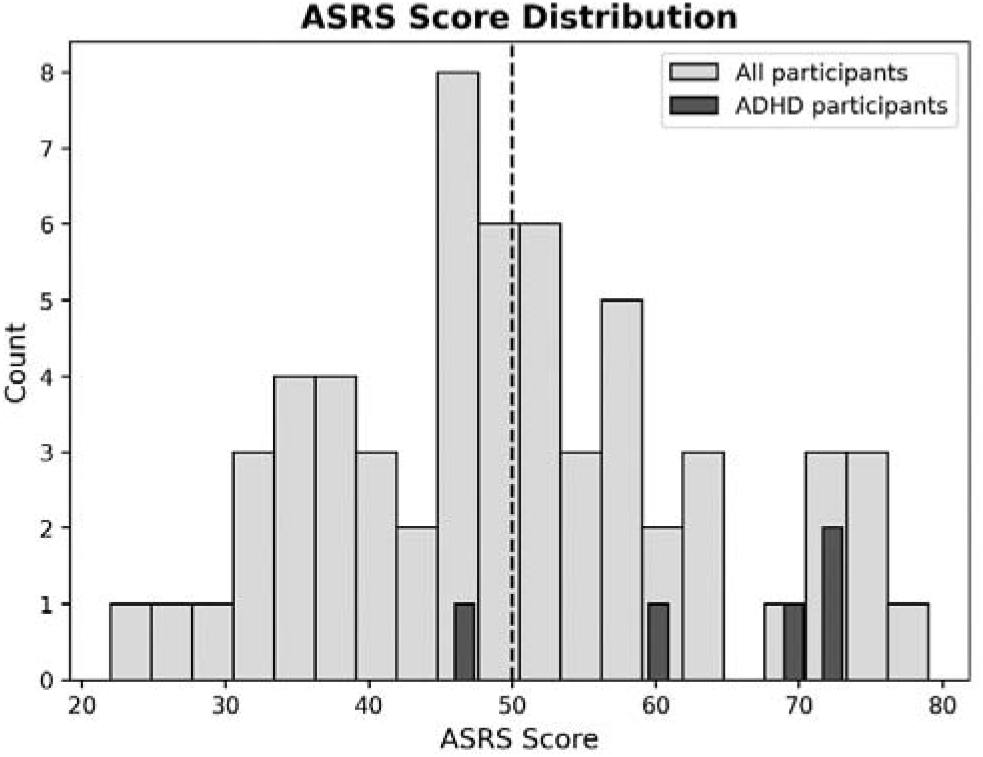
ASRS Scores. Histogram showing the distribution of ASRS scores across participants without ADHD (light gray) and with ADHD (dark gray). ASRS scores were widely distributed across the sample, with a mean score of 50 (SD = 13.21).

## Discussion

The goal of this study was to evaluate different behavioral and neural metrics that are associated with attentional functioning and have been proposed as reliable predictors of individual differences. The current results are sobering in this regard and, in line with several recent meta-analyses (Dean et al., 2025a; Eisenberg et al., 2019; Mazza et al., 2021a), show a lack of convergence between most behavioral measures commonly used to index attentional capabilities. Specifically, in this dataset we found no correlation between behavioral metrics associated with executive control and phasic alertness (derived from the ANT task), working-memory capacity (derived from the N-Back task) and fluid intelligence (derived from the Ravens’ APM task). Moreover, none of these behavioral metrics predicted self-reported symptoms of ADHD (derived from the ASRS questionnaire), raising further questions regarding their ability to account for difficulties in attention as experienced in daily life (Du Rietz et al., 2016; Toplak et al., 2013b).

The EEG-based neural measures of ISC were somewhat more promising in their ability to capture consistent differences between individuals. ISC measures showed consistent convergence both within paradigm (ISC to different stimuli) and across paradigms (Auditory Oddball and Attention to Speech tasks). This suggests that the degree of similarity/divergence of an individual’s neural activity relative to the group is stable, across experimental contexts. Although ISC-stability in and of itself is not a direct indicator of attention, in the Attention to Speech task we did find that Continuous-ISC was significantly and positively correlated with comprehension accuracy. This aligns with proposed interpretations of ISC as reflecting one’s level of engagement with stimuli presented to them (Dmochowski et al., 2012, 2014; Ki et al., 2016b; Rosenkranz et al., 2021), and supports its possible utility for capturing stable individual differences in attention-related processes.

### The promise of ISC and how to interpret it

The notion of inter-subject correlation (ISC) was originally introduced in fMRI studies (Hasson et al., 2004), and later adapted to EEG (Dmochowski et al., 2012; Nastase et al., 2019), as a way to quantify the similarity of neural responses across individuals exposed to continuous stimuli. In contrast to traditional event-based approaches, that examine neural responses to discrete well-defined events, ISC is particularly useful for studying complex naturalistic stimuli, such as movies, stories, or lectures, in which information unfolds continuously over time. The basic premise is that common temporal dynamics across participants can be interpreted as driven by external stimuli, whereas neural activity related to internally generated thoughts or other idiosyncratic activity leads to less similarity between participants (Nastase et al., 2019). Following this logic, modulation of ISC has been linked to ones’ level of attention or engagement with a presented stimulus: participants who are more engaged show neural responses that are more similar among themselves (driven by the joint stimulus), whereas individuals who are more distracted and don’t follow the stimulus as closely, would exhibit reduced ISC (Ki et al., 2016a; Madsen & Parra, 2022; Rosenkranz et al., 2021).

In the present study, we estimated ISC in two separate paradigms and assessed ISC both in response to specific events (ERP-ISC) and continuously (Continuous-ISC). Results showed convergent patterns, both within and across paradigms: ERP-ISC to the three stimuli in the Oddball paradigm (Standard, Target, and Novel) were highly correlated with each other, as were ERP-ISC to disruptions and the Continuous-ISC in the Attention to Speech task. Moreover, across paradigms, ERP-ISC to disruptions in the Attention to Speech task was correlated with ERP-ISC to Target and Novel stimuli in the Oddball paradigm, and Continuous-ISC was correlated ERP-ISC to Novel sounds. This pattern is consistent with the possibility that the idiosyncrasy of ones’ neural responses, in this case to auditory stimuli, is stable for both highly controlled and more ecological contexts. These findings also align with a previous study from our group, where we demonstrate stable within-subject test-retest reliability of ERP-ISC in the Oddball paradigm (Korisky et al., 2026).

Following the rationale linking higher ISC with higher engagement with the task and better stimulus processing, these findings may suggest that ISC effectively captures stable within-person tendencies to maintain attention to a task at hand (Finn et al., 2020; Gao et al., 2020). Supporting this interpretation, previous studies have shown that ISC to continuous stimuli tracks selective attention in competing-speaker paradigms, decreases when attention is diverted away and increases during emotionally salient and narratively meaningful moments (Dmochowski et al., 2012; Ki et al., 2016a; Madsen & Parra, 2022; Nastase et al., 2019; Rosenkranz et al., 2021). Other studies further show that ISC is positively correlated with learning outcomes, attentional engagement (with the shared stimulus), and memory performance (Cohen et al., 2018b; Cohen & Parra, 2016; Madsen & Parra, 2022), across both naturalistic and semi-controlled paradigms (Liu et al., 2025). This interpretation aligns with the positive correlation found here between performance on the Attention to Speech task and Continuous-ISC.

And yet, several caveats persist. First, with the exception of the positive association between Continuous-ISC and performance in the Attention to Speech task, variations in ISC were not consistently correlated with the broad set of cognitive-behavioral measures collected here, including self-reported ADHD symptoms, leaving a crucial gap in our ability to explain the factors underlying these individual differences. Second, dissimilarity in neural responses across individuals can arise from multiple sources other than differences in engagement. For example, reduced ISC may arise from variation in the time-course of neural responses, anatomical differences that alter the spatio-temporal projection of neural sources at the scalp, or non-neural contributions such as eye movements and other physiological signals (Cohen et al., 2018b; Ki et al., 2016a; Madsen & Parra, 2022; Nastase et al., 2019; Nielsen et al., 2023; Poulsen et al., 2017; Srinivasan, 2006). Similarly, a variety of other personal or cognitive factors, not necessarily related to engagement/attention have been shown to affect neural response idiosyncrasy, including social impairments, emotional regulation, personality traits and reading abilities (Finn et al., 2018, 2020; Jangraw et al., 2023; Matz et al., 2022; Ramot et al., 2020). Given this, despite the promise of ISC to provide an objective, data-driven indication of participant engagement/attention to continuous stimuli, and the nice convergences found here, additional research and perhaps new tools are needed to separate the contributions to ISC stemming from true variation in cognitive processing of the stimuli from additional task non-specific variance.

### Generalization across artificial and ecological contexts

One of the main goals of this cross-paradigm study was to test which aspects of neural activity generalize between artificial and semi-ecological approaches for studying auditory attention. Besides the correlations in ISC discussed above, perhaps the most striking effect was the strong resemblance between ERPs to Novel sounds in the Oddball task and to the onset of disruption events in the Attention to Speech task. These stimuli differ from each other in many ways – from the properties of the *stimuli* themselves (short ecological sounds vs. speech-utterances), to the *context* in which they are presented (Novel sounds in the Oddball task are part of the to-be-attended sequence whereas Disturbances in the Attention to Speech task were presented from behind, concurrent with target audio-visual speech); and the specific *task* demands (Novel sounds in the Oddball task need to be discriminated as non-target whereas Disturbances in the Attention to Speech task need simply to be ignored, while attempting to comprehend the video lesson). Despite these vast differences, from an attention perspective, in both cases, these stimuli constitute salient unexpected events that are hypothesized to capture bottom-up attention (Escera et al., 1998; Escera & Corral, 2008; Polich, 2007). And indeed, the ERPs generated for both stimuli were highly similar and contained a sequence of components associated with perceptual salience and the capture of attention. These included an enlarged N1 response, followed by a prominent P3-like response and a later negative component, all of which shared a common centro-frontal scalp distribution characteristic of auditory responses.

The N1 component captures early sensory responses, generated in auditory cortex, which are considered mandatory bottom-up responses to external stimuli (Jääskeläinen et al., 2004; Näätänen & Picton, 1987). However, later ERP responses are usually associated with higher-level cognitive processes, pertaining to the interpretation and relevance of specific stimuli, beyond their basic acoustic features. As such, the family of P3 responses are hallmarks of surprise-related processes, associated with attentional capture/re-orienting, stimulus evaluation, target detection, and context updating, with some distinction between task-relevant targets and novel, unexpected sounds (Escera & Corral, 2007; D. Friedman et al., 2001; Polich, 2007). In this respect, there is a functional parallel between Novel sounds in the Oddball task and disturbances in the Attention to Speech task, which both constitute unexpected and salient non-target events that may elicit involuntary orienting-related responses, despite (and perhaps because of) their task-irrelevance (Berti, 2008, 2013; Fiedler et al., 2025; Levy, Libman Hackmon, et al., 2025; Wetzel et al., 2013). Similarly, the later negative component, which peaked at around 400 ms is reminiscent of the N400 response typically found in language studies for unexpected words and is associated with semantic surprisal (Friederici et al., 1993a; Holcomb & Neville, 2013a; Kutas & Federmeier, 2011a). Although the current experiment was not explicitly designed to manipulate semantic surprisal, both the novel sounds in the Oddball task and the disruptions in the Attention to Speech task carry clear semantic value, that may underlie later response (Brown et al., 2023; Friederici et al., 1993b; Holcomb & Neville, 2013b; Kutas & Federmeier, 2011b).

The observed similarity in neural responses across the two tasks – in both ERPs and ISC - is extremely encouraging, given ongoing efforts to increase the ecological validity of attention research (Hölle et al., 2021; Korte et al., 2025; Levy, Korisky, et al., 2025; Levy, Libman Hackmon, et al., 2025; Liebherr et al., 2021). Finding that similar neural responses are generated in highly controlled artificial tasks and under more naturalistic conditions contributes to establishing their generalized validity, on the continuum between the lab and real-life (Janssen et al., 2021).

### The challenge of cognitive-behavioral measures for evaluating individual differences in attention

As opposed to the neural measures that showed some convergence across paradigms, the behavioral-cognitive measures presented a less consistent picture. Given the extensive literature on the ANT task, and its proposed utility for assessing different types of attention-related processes, we were surprised to find that none of the behavioral measures – overall accuracy, RT, or their modulation by Cognitive Control demands (flanker congruency) and Alertness (a preceding tone) – were correlated with each other within subject, nor did they predict any of the neural measures or ADHD symptoms. This stands in contrast to its proposed utility for exposing differences in attention abilities, especially in clinical populations such as ADHD (Cao et al., 2008; Johnson et al., 2008; Lambek et al., 2011) and Schizophrenia (Arora et al., 2020; Bieleninik et al., 2023; Hahn et al., 2011). However, test-retest analyses of ANT score reliability have shown only modest within-person stability (Ishigami & Klein, 2010b; MacLeod et al., 2010b), suggesting that ultimately it is less useful for capturing individual trait-based differences (Hahn et al., 2011; Ishigami & Klein, 2010a; Kong et al., 2024; MacLeod et al., 2010a; Paap & Oliver, 2016).

The battery of tasks used here also included two tasks that evaluate more general cognitive abilities – namely, working memory and fluid intelligence – that are often considered to be important for attentional functioning (Tsukahara et al., 2025; Zhao & Vogel, 2025). Neuroimaging studies have shown that these supposedly distinct cognitive constructs often rely on shared fronto-parietal networks, leading to suggestions that they may, in-fact, capture more domain general attributes (Assem et al., 2020; Corbetta et al., 2008; Fedorenko et al., 2013; Santarnecchi et al., 2021; Seeburger et al., 2026). Given this, here too we had expected to find that scores on the N-back and Raven’s APM tasks, which are highly-accepted operationalization for measuring these constructs, would explain at least some individual-variance on the attention-related measures collected here. However, this was not the case, as neither measure correlated with ANT-outcomes or any of the neural measures from the Oddball or Attention to Speech tasks. Moreover, scores on the n-back and Ravens tasks were only loosely correlated with each other (and did not survive correction for multiple comparisons), contributing to ongoing discussion regarding the relationship between them (Ackerman et al., 2005; Burgoyne et al., 2025; Kane et al., 2007; Redick & Lindsey, 2013). Adding to that, in the current dataset ASRS scores, which capture one’s introspective perception of their own attentional difficulties and are used clinically to evaluate the severity of ADHD symptoms, also did not correlate with any of the other measures collected – behavioral or neural.

Hence, the emerging picture is of a marked lack of convergence across different empirical metrics that are all hypothesized to capture at least partially-overlapping aspects of the construct “attention”. This pattern is not unique to the present study. With the rising emphasis on replicability and recognition of the prevalence of underpowered studies in cognitive psychology and neuroscience (Baker et al., 2021; Button et al., 2013; Szucs & Ioannidis, 2017), we see an increase in large meta-analyses and multi-task-battery studies exposing the poor convergence of data across supposedly similar experiments. For example, (Mazza et al., 2021) found that across 37 cognitive tasks and 138 task-derived variables, the mean absolute correlation between survey and task measures of self-regulation was only .049. Similarly, (Eisenberg et al., 2019) reported that task-based and self-report measures of self-regulation showed little empirical overlap and did not resolve into a single coherent construct, and Dean et al., (2025b) showed that several experimental measures were so weakly related to the rest of the battery that they had to be excluded from factor analysis, including interference scores derived from the ANT, Stroop, and task-switching paradigms. Similar inconsistencies are found when focusing specifically on measures relating to attention. For example, factor-analytic evidence suggests that putative attention tests do not form a single unified factor, but instead fractionate into separable components (Treviño et al., 2021). Similarly, classic measures of executive control such as Flanker, Stroop, and Simon-type effects exhibit poor convergent validity across tasks (Paap & Oliver, 2016; Whitehead et al., 2020) and Continuous Performance Tasks (CPTs), which are among the most common tools for assessing difficulties in sustained attention, also have been shown to carry only modest-to-moderate standalone-sensitivity to capture individual differences in attention (Arrondo et al., 2024b; Shaked et al., 2020). Accordingly, the mostly null-correlations found here unfortunately fits with the broader confusion as to the mapping of complex cognitive constructs onto task-specific operationalizations and behavioral measures.

Several explanations are commonly given to address this lack of convergence, from methodological and measurement-related concerns such as sample size, signal noise and multiple comparisons (Perneger, 1998), through acknowledging the limitations of computerized tasks (Webb & Demeyere, 2023; Yangüez et al., 2023) to more theoretical accounts pertaining to the dissociable nature of sub-processes related to the broader construct of “attention” (Fuermaier et al., 2015; Toplak et al., 2013b). Most likely, ***all*** of these factors play some role in the current difficulty to identify simple, consistent and highly replicable cognitive-behavioral measures to quantify “attention”. However, to date, this operational challenge and conceptual ambiguity poses a significant barrier for scientific progress in understanding the neurobiological and behavioral substrate underlying attention-functioning, in real-life.

### Toward an integrated framework for measuring attention across contexts

The present work makes several important contributions toward overcoming this barrier and laying a path towards an integrated framework for studying attention. First, it emphasizes **convergence and generalizability** as a central criterion for validating proposed measures, which should serve as a principled approach toward narrowing-down the number of candidate-metrics for quantifying “attention” out of the numerous candidates used in current scientific and clinical work (Burgoyne et al., 2025; Draheim et al., 2022b; Hedge et al., 2017). Second, our work specifically focuses on convergence between artificial and ecological tasks, which is critical for **bridging the gap between lab-testing and the real-life challenges of attention** (Janssen et al., 2021; Levy, Libman Hackmon, et al., 2025). By showing that similar neural responses are generated for surprising events across both the oddball and attention to speech paradigms used here, the current results provide an important proof of concept of the possibility to bridge this gap and identify neural measures that are relevant to real-life contexts. Last, the convergence between ISC measures observed here, both within and between tasks, supports the idea the **idiosyncrasy in the pattern of an individuals’ neural activity** can provide a promising and generalizable metric of engagement and/or attention, with the potential to overcome the invariable task-specific differences across contexts and ultimately to extend the ecological validity of attention research.

## Acknowledgments

This work was funded by the Israel Science Foundation, grant # 274/24 to ezg.

